# *In situ* architecture of the basal body reveals regionalized assembly and central pair microtubules nucleation

**DOI:** 10.64898/2026.10.08.757802

**Authors:** Ziyan Wang, Huan Yang, Hao Lang, Xueliang Zhu, Yun Zhu, Fei Sun

## Abstract

The basal body (BB) is essential for axoneme assembly and ciliary function, and its defects are associated with ciliogenesis disorders and ciliopathies^1,2^. However, the molecular basis of the BB’s region-specific transitions along the proximal-distal axis, and the mechanism by which ciliary central pair (CP) microtubules are nucleated during ciliogenesis, have remained largely unknown. Here, we resolve the subnanometer architecture of native BBs in mammalian multiciliated cells using cryo-electron tomography and subtomogram averaging, and identify 11 ciliary-base microtubule-associated proteins (CMAPs), including 5 previously uncharacterized core components. These CMAPs localize specifically along ciliary base that facilitate the triplet-to-doublet microtubule transition. We further identify luminal γ-tubulin ring complexes (γ-TuRCs) anchored to the microtubule wall via the augmin complex, and propose that they may serve as candidate nucleation sites for CP microtubules biogenesis. Together, these findings define a molecular blueprint of the BB and provide a structural framework for understanding how regional architectural specialization and luminal nucleation collectively drive ciliary assembly and how their dysfunction causes ciliopathies.

## Main

Basal bodies (BBs) are centriole-derived organelles that template ciliary assembly, and support cilia-mediated motility and sensory signalling across eukaryotes^3–5^. By defining the ninefold symmetry of the axoneme and coordinating membrane docking, microtubule organization and cellular polarity, BBs establish the structural framework for ciliary function^6^. The role is particularly critical in motile cilia, where the ordered arrangement and coordinated orientation of BBs provide the structural basis for synchronized ciliary beating. However, the architectural complexity and native organization of BBs has long posed a major challenge to high-resolution imaging and molecular dissection in situ.

The transition of the BB into the ciliary axoneme raises two long-standing questions in the field. First, as the microtubule triplet (MTT) structure of the BB is progressively remodelled into the doublet configuration of the axoneme, how the structural integrity is maintained at each transitional intermediate remains unknown^7^. Second, the mechanism by which the ciliary central pair (CP) microtubules are nucleated de novo during ciliogenesis is unresolved^8^. Although genetic and cell-biological studies have implicated luminal factors near the transition zone (TZ) and basal plate^9–11^, direct in situ structural evidence for CP microtubule initiation is still lacking.

Over the past 10 years, ultra-expansion microscopy (U-ExM), cryo-electron microscopy (cryo-EM) and cryo-electron tomography (cryo-ET) have greatly advanced our understanding of centrioles and BBs to the molecular level^12–16^, revealing multiple subregional components, including A-C linker, microtubule inner proteins (MIPs); and luminal ring factors. However, the native architecture of the BB and its direct contribution to ciliary motility remains incompletely understood, because in situ approaches have lacked sufficient resolution, and isolated preparations often disrupt native organization^17,18^. Here, we present a subnanometer-resolution in situ structure of the native BB in mammalian multiciliated cells (MCCs). We identify 11 ciliary-base microtubule-associated proteins (CMAPs) distributed along the proximal–distal axis, revealing the molecular basis for stabilizing the triplet-to-doublet transition. In addition, we determine the in situ architecture of luminal γ-tubulin ring complexes (γ-TuRCs) complexes anchored to the microtubule wall via the augmin complex, and provide structural evidence linking luminal nucleation machinery to CP microtubules initiation. Together, these findings define a comprehensive molecular blueprint of the BB and establish a unifying structural framework for understanding ciliary assembly and its disruption in disease.

## Results

### Overall architectures along the ciliary base

We acquired cryo-ET datasets of BBs from mouse ependymal cells (mEPCs) (**Fig. S1**) and revealed a highly ordered, region-specific architecture along the proximal-distal axis (**Fig. 1a and Video S1**). The pinhead, inner scaffold and luminal ring define a stepwise reinforcement framework that stabilizes distinct regions of the BB along this axis (**Fig. 1b**). We also observed luminal γ-TuRCs, which display a distribution pattern overlapping with that of the inner scaffold (**Fig. 1b**), indicating a potential spatial association.

**Figure 1.**
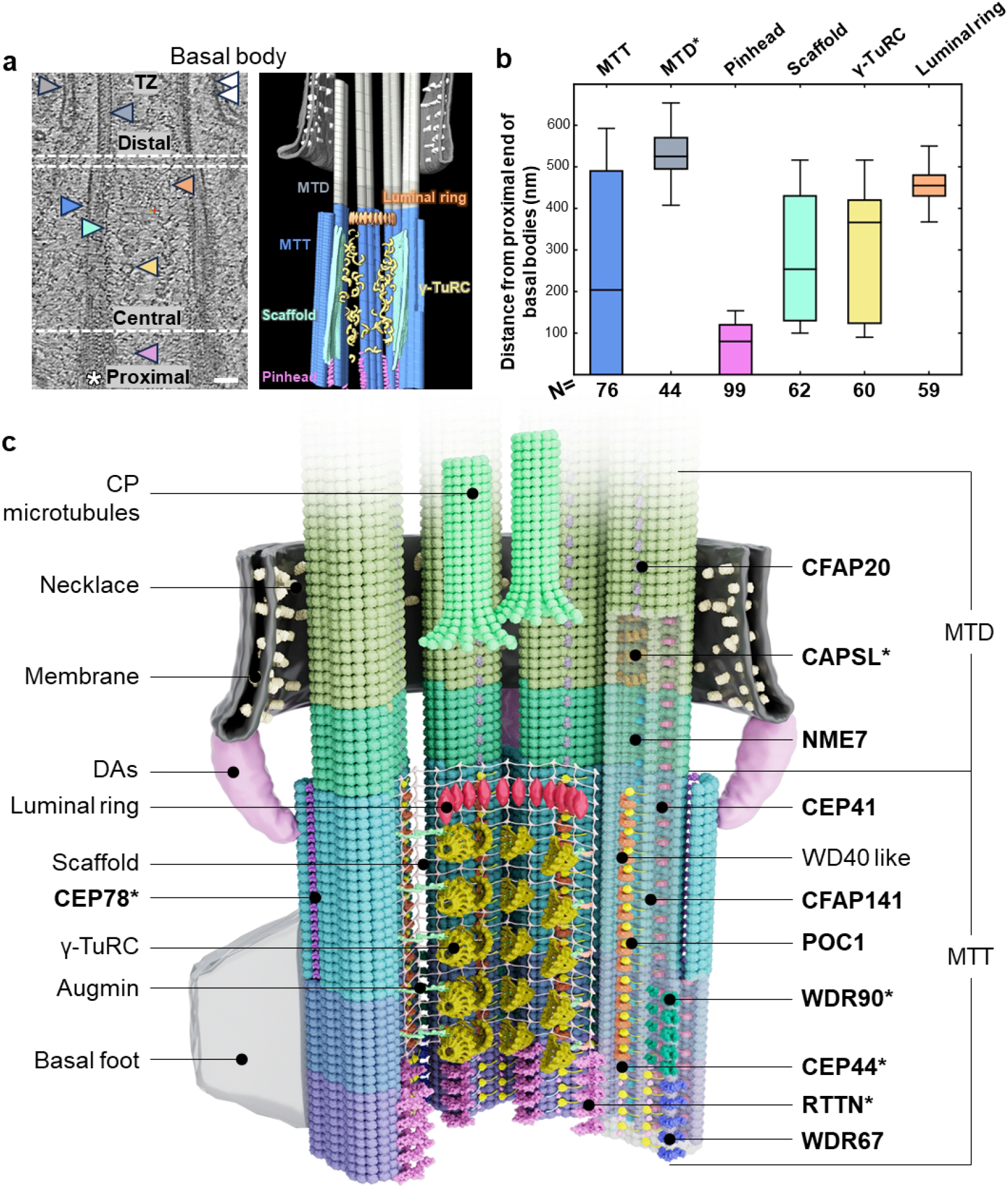
Architecture of mammalian BB. **(a)** 2D slice through a tomogram of mouse mEPCs, showing a ciliary base in longitudinal cross section (left panel). A 3D rendering of the tomogram corresponding to the left panel is shown on the right panel. The major components are indicated and labelled. Scale bar: 50 nm. Blue: MTT/MTDs; orange: luminal ring; purple: pinhead; green: inner scaffold; yellow: luminal γ-TuRC; grey: ciliary pocket; white: necklace. **(b)** Position along the ciliary base occupied by different structures. The x-axis represents structural components and the corresponding number of statistical substructures, and the y-axis represents their distributions, measured relative to the bottom of BB (marked by the white asterisk in panel a). MTD* denotes the segment of the MTD up to the point where the membrane appears. Only high-quality tomograms in which the asterisk-marked reference was clearly visible were analysed. For each fully resolved substructure, its axial range was quantified as the distance from the asterisk reference to the positions where the density first appeared and disappeared. **(c)** Model of a BB displaying the structural elements along its proximal end to TZ. CMAPs are shown in bold; newly identified proteins are marked with asterisks.

Specifically, we obtained five distinct structural types of MTTs and microtubule doublets (MTDs) by subtomogram averaging (STA), with local resolutions up to 5.3 Å (**Fig. S2-S3 and Table S1**), enabling integrated structural modelling as reported in our previous study^19^. By integrating mass spectrometry (MS) (**Data S1-S3**), AlphaFold2^20^, FindMySequence^21^ and DomainFit^22^, we unambiguously assigned tubulin isoforms and 11 distinct CMAPs into these maps (**Fig. 1c, Fig. S4-S10, Table S2-S3 and Video S2**). Notably, nearly all assigned components are supported by independent protein–protein interaction (PPI) predictions^23^, providing cross-validation for our molecular assignments (**Fig. S11**). Together, these results establish an annotated, region-specific in situ structural framework for the mammalian ciliary base (**Fig. 1c**).

This pronounced regional heterogeneity in molecular composition underscores the structural and functional complexity of BBs. Region-specific CMAPs serve as molecular markers that demarcate functional distinct BB regions, shedding light on the multifaceted roles of BBs—ranging from template-mediated duplication at the proximal end to membrane docking and ciliary axoneme biogenesis at distal sites.

### Regional CMAP patterning stabilize microtubule transition of the ciliary base

The MTT architecture of the BB is progressively remodelled into the MTD configuration of the ciliary axoneme, and this transition is accompanied by spatially ordered remodelling of CMAPs, rather than uniform distribution of stabilizing factors.

The proximal region of BB harbours the most specialized structural modules: the pinhead links cartwheel-associated structure to microtubule wall, and the A-C linker interconnects adjacent MTTs (**Fig. S12a**). Here, we identify RTTN as the core component of the pinhead (**Fig. 2a-b, Fig. S5 and Fig. S12**). RTTN forms an elongated ARM-repeat-containing density, whose C-terminal region contacts protofilament A04 via electrostatic interactions, thereby coupling internal proximal structures to the A-tubule wall (**Fig. 2c and Fig. S5**). In addition, a WD40-like density is present in the pinhead of mature BBs but absent in immature BBs and lower species (**Fig. S12b-d**), suggesting that this component is recruited post-maturation to further reinforce the proximal BB. In the A-C linker, WDR67 forms a major part of the C-link through extensive contacts with the C-tubule (**Fig. S7**), as previously reported^12,15^. Guided by AlphaFold2-assisted fitting into the corresponding A-link densities, we propose WRAP73 and CEP135 as candidate components constituting the mammalian A-C linker (**Fig. S13**). Together, the pinhead and A-C linker establish a proximal reinforcement module that couples the cartwheel–pinhead axis to inter-triplet cohesion.

**Figure 2.**
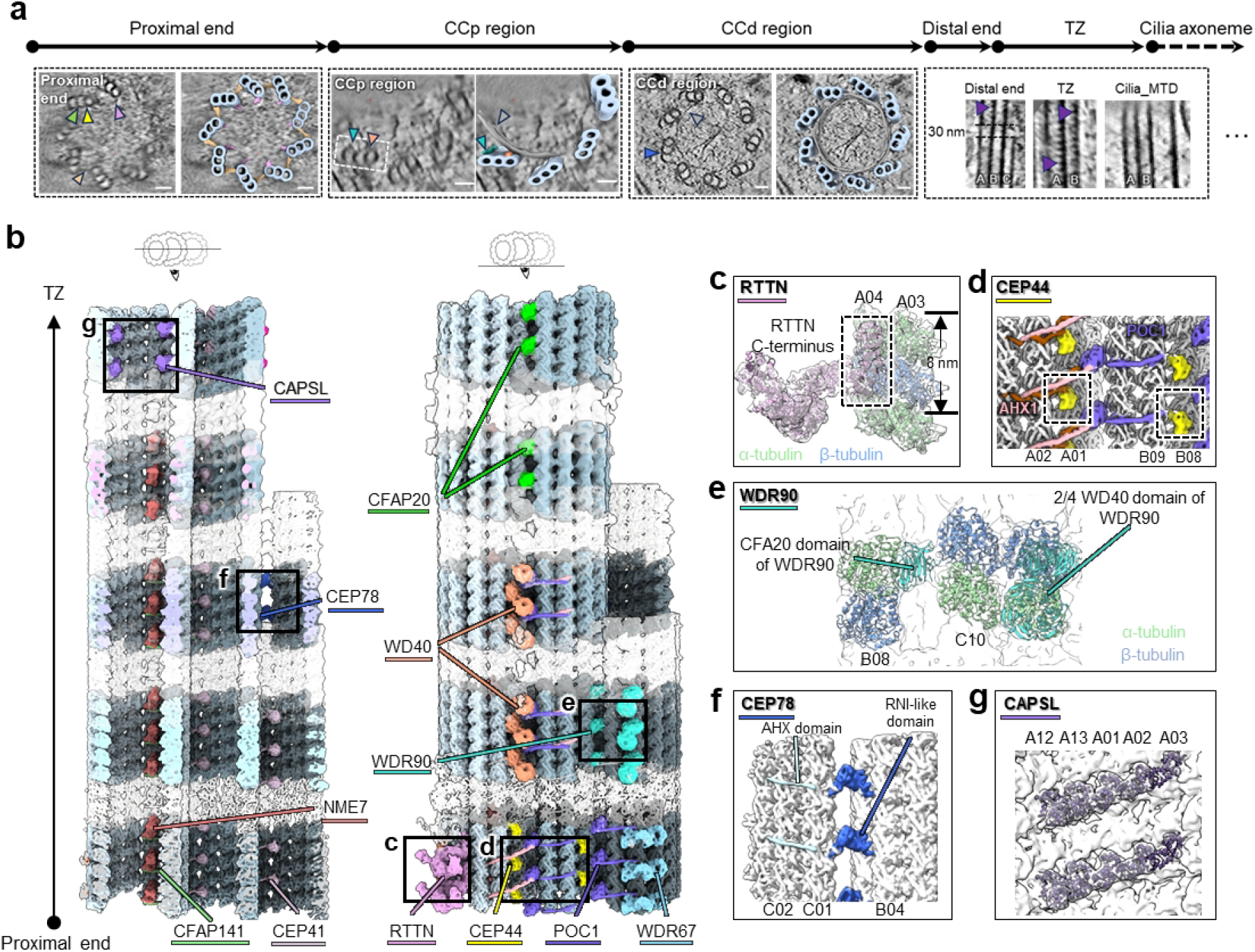
In situ structures and region-specific distributions of CMAPs along the ciliary base. (a) Representative tomogram slices of the 5 ciliary base regions in top or side views. The key structures are labelled in arrowheads. Light blue: MTT/MTD; purple: pinhead; yellow: A-B inner junction; green: B-C inner junction; grey: inner scaffold; Dark blue: B-C out junction. Scale bar: 50 nm. (b) Composite in situ structures of the 5 ciliary base regions, aligned along the proximal-distal axis to illustrate spatial relationships, with transparent densities showing structural transitions between adjacent regions. The CMAPs are indicated and coloured differently. (c)-(g) Magnified views of key CMAPs and their molecular interactions in the corresponding regions marked by black boxes in (b).

POC1 has been established as a core component at the A-B and B-C inner junctions of BBs^16,24^. At the proximal BB, we further identify CEP44 as an additional component occupying both the A-B and B-C inner-junctions, in contrast to prior reports placing it exclusively in the pinhead^13^ (**Fig. 2d, Fig. S6 and Fig. S12a**). At the A-B inner junction, CEP44 binds directly to protofilament A01, with an additional helical density-likely contributed by CEP44-further stabilizes this interaction (**Fig. 2d**). These observations indicate that CEP44 and POC1 act cooperatively to reinforce the A-B interface and preserve structural integrity of the proximal BB.

As the structure transitions into the central core region, the cartwheel and A-C linker disappear, accompanied by the emergence of an inner scaffold for continuous internal support and a gradual transformation of the full C-tubule into a half C-tubule (**Fig. 1**). Correspondingly, RTTN, CEP44 and WDR67 are no longer present, whereas new CMAPs gradually assemble (**Fig. 2b**). POC1 remains a key continuous component at the A-B inner junction, while four WD40-like densities emerge adjacent to each POC1 molecule (**Fig. S14a-c**), consistent with previous study^14^. Focused STA further reveals a hook-like scaffold feature protruding toward the lumen (**Fig. S14d**), indicating a potential interface between the scaffold and luminal components in the BB central core.

In the <u>c</u>entral <u>c</u>ore region adjacent to the <u>p</u>roximal BB (hereafter termed the CCp region), the B-C inner junction exhibits pronounced regional specialization coinciding with retention of the full C-tubule (**Fig. 2a and Fig. S14a**). We identify WDR90 at the B-C inner junction of the CCp region via its CFA20 domain (**Fig. 2e and Fig. S8**), where it provides structural support for this interface. In the <u>c</u>entral <u>c</u>ore region adjacent to the <u>d</u>istal BB (hereafter termed the CCd region), another WD40-like density is observed exclusively attached to protofilaments B07-B08 (**Fig. 2a and Fig. S14b**), leaving the B-C outer junction as the main connector for the C-tubule. Here, we identify CEP78 as the core bridging component (**Fig. 2f and Fig. S9**). The RNI-like domain of CEP78 engages both the B- and C-tubules while an extended α-helix wraps around three C-tubule protofilaments (**Fig. 2f**). This microtubule-binding mode effectively stabilizes the half C-tubule in the absence of a canonical B-C inner junction.

At the distal BB, MTTs terminate and transition into MTDs within a short axial distance (∼30 nm), after which the TZ extends to the site of ciliary CP microtubules initiation (**Fig. 2a**). NME7 and CEP41 persisted as microtubule inner proteins (MIPs) from the proximal region to the distal BB (**Fig. 2b**), consistent with previous reports^14^. By contrast, CFAP20 emerged at the A-B inner junction of MTDs to replace POC1, indicating that the A-B junction has begun adopting axonemal characteristics (**Fig. S15a-b**). Within the TZ, a distinct set of MIP densities emerged in A- and B-tubules, forming a semicircular belt around the tubule wall that provides strong stabilization (**Fig. 2g**). This density centres on α-tubulin, rather than occupying grooves between α- and β-tubulin subunits as reported for SPACA9^19,25^. Following screening of tubulin-interacting proteins and DomainFit analyses, CAPSL is assigned to this density (**Fig. S10**), further supported by its detection in our MS list of isolated BB clusters and immunofluorescence (IF) localization (**Fig. S16 and Data S3**).

Collectively, these data reveal proximal-to-distal sequential remodelling of CMAPs from the BBs to ciliary axoneme: proximal RTTN, CEP44, WDR67 couple the cartwheel-pinhead axis to triplet cohesion; WDR90 and CEP78 selectively stabilize the full C-tubule and half C-tubule in the central core region, respectively; and distal/TZ localized CFAP20 and CAPSL mark the structural transition toward axonemal identity.

### CMAPs play distinct role in BB integrity and ciliary motility

To directly investigate the functional contributions of newly identified CMAPs in the BB, we performed RNA interference (RNAi)-mediated knockdown in mEPCs and assessed consequent impacts on BB structural integrity and motile ciliogenesis (**Fig. 3, Fig. S17-S19 and Video S3-S7**).

**Figure 3.**
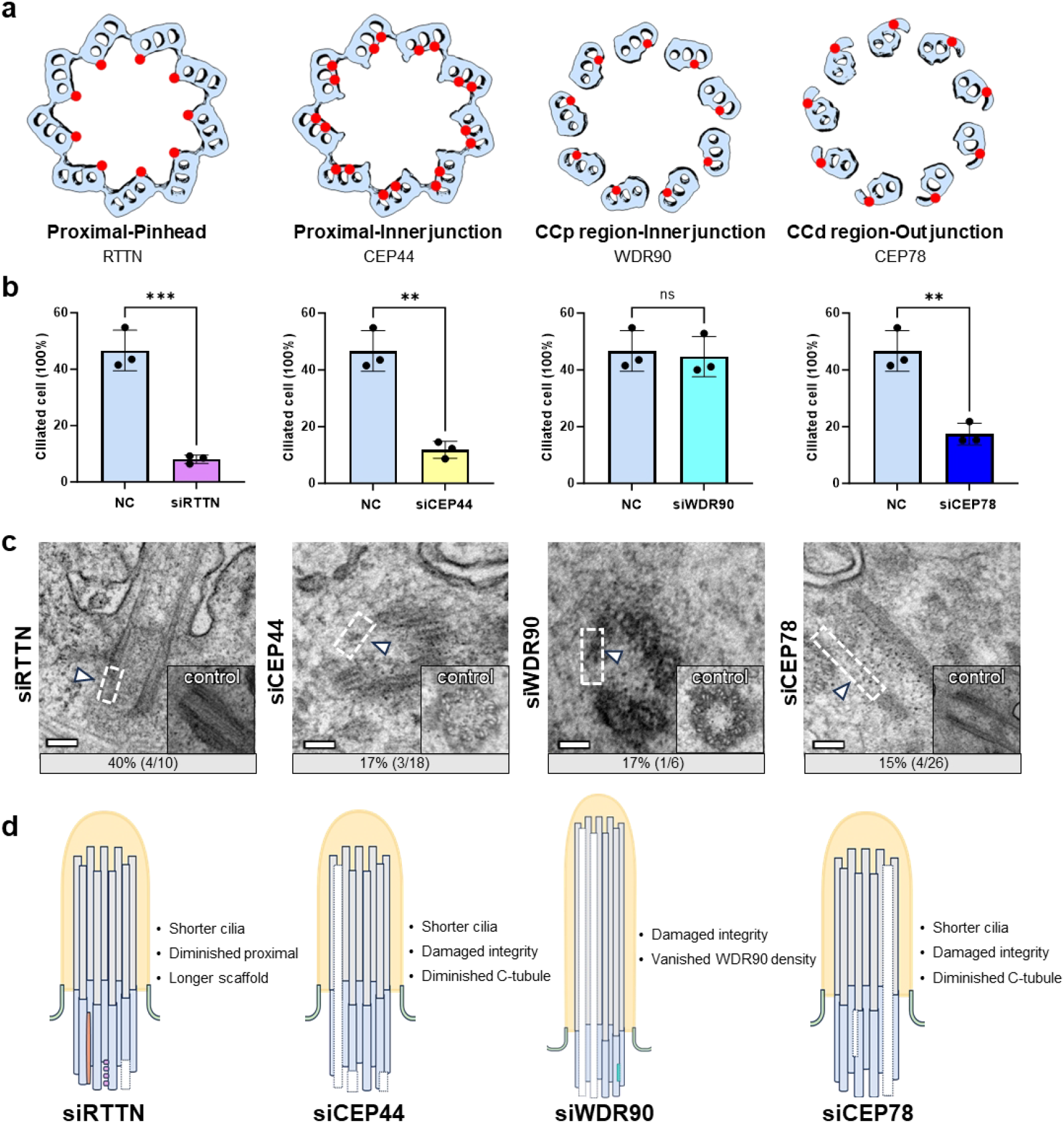
RNAi-mediated depletion of key CMAPs disrupts BB architecture and ciliary function in mEPCs. **(a)** Schematic of the spatial localization of four critical CMAPs in BB. **(b)** CMAPs depletion repressed multiciliogenesis in different degree. The data are from three independent experiments. For each experiment, n > 500. Statistical significance was determined using a t-test. **(c)** TEM images of BB ultrastructure in CMAP-depleted Day 10 mEPCs. Scale bar: 100 nm. **(d)** Schematic summary of BB and ciliary phenotypes upon CMAPs depletion.

As the predominant CMAP of the pinhead, RTTN likely acts as a circumferential “reinforcement node” that mechanically stabilizes the centriole and BB. Consistently, RTTN depletion markedly reduced the percentage of ciliated cells and shortened the length of cilia, with most remaining cilia exhibiting rotational movement (**Fig. 3b and Fig. S17-S19**). Conventional transmission electron microscopy (TEM) revealed that some MTTs were absent and that the inner scaffold replaced the pinhead in some proximal MTTs (**Fig. 3c**). The cryo-ET data further confirmed this variation (**Fig. S19b**). These observations not only support our structural assignment of RTTN as a pinhead component, but also uncover its essential function in sustaining BB architecture and enabling normal ciliary motility.

In the same region, CEP44 connected the A-B and B-C tubules in collaboration with POC1, thereby providing structural support for the proximal BB. Consistent with RTTN depletion, CEP44 depletion also caused a significant reduction in cilia length and ciliation efficiency (**Fig. S17-S19**). Moreover, TEM and cryo-ET analyses revealed that the completeness of the proximal MTTs in the BB was disrupted (**Fig. 3c and Fig. S19b**). Further STA data revealed structural variations in the BB upon CEP44 depletion (**Fig. S19c**). Rather than a global collapse, we observed a preferential loss of density at the B-C junction, whereas the A-B inner junction remained largely intact (**Fig. S19c**). This structural preservation suggests that CEP44 exhibits higher affinity and stability at the A-B junction, indicating that this is the primary anchor point critical for maintaining BB integrity.

As the connector of B-C inner junction in CCp region, WDR90 appears to enhance the stability of MTT. Depletion of WDR90 did not cause significant defects in ciliary length or motility compared with the negative control (**Fig. S17-S19**). However, TEM and cryo-ET data revealed that the BB architecture was severely compromised in the siWDR90 group (**Fig. 3c and Fig. S19b**), consistent with previous work^26^. STA map further demonstrated a specific loss of densities along the C-tubule in the seemingly preserved MTT CCp region, indicating that WDR90 is essential for the assembly and stabilization of the C-tubule (**Fig. S19d**).

The structure of CEP78 in the CCd region strongly supports its role as a connector between the half C-tubule and the adjacent B-tubule. CEP78 depletion significantly reduced the ciliogenesis rate and cilia length, triggered rotational movement in ∼35% of cilia, and induced severe structural damage to the BBs (**Fig. S17-19 and Fig. 3**). Specifically, MTTs with half-C were missing, whereas MTDs continued to extend. In many cases, MTTs were completely absent, leaving only single microtubules derived from MTDs. These defects indicate that CEP78 is critical for maintaining the proper structural organization of BBs and cilia.

Collectively, these functional analyses establish that region-specific CMAPs are not merely structural components but master regulators of BB ninefold symmetry, ciliary assembly, and motile function. Their deficiency simultaneously disrupts BB structural integrity, reduces ciliary length, and distorts ciliary beating patterns, revealing a direct mechanistic link between BB architectural stability and ciliary functional output.

### The luminal ring-associated structure mediates BB docking to the plasma membrane

Previous work identified a specialized a luminal-ring with 27-fold symmetry at the central-to-distal transition of BBs and centrioles, yet its functional role remained unclear^27–29^. Our *in situ* cryo-ET analysis revealed that luminal-ring associated density connected to the distal appendages (DAs) and extended to anchor at the cell membrane (**Fig. 4a-b and Fig. S20**). We propose that the luminal ring forms a continuous structural linkage that participates in the docking of BBs to the cell membrane.

**Figure 4.**
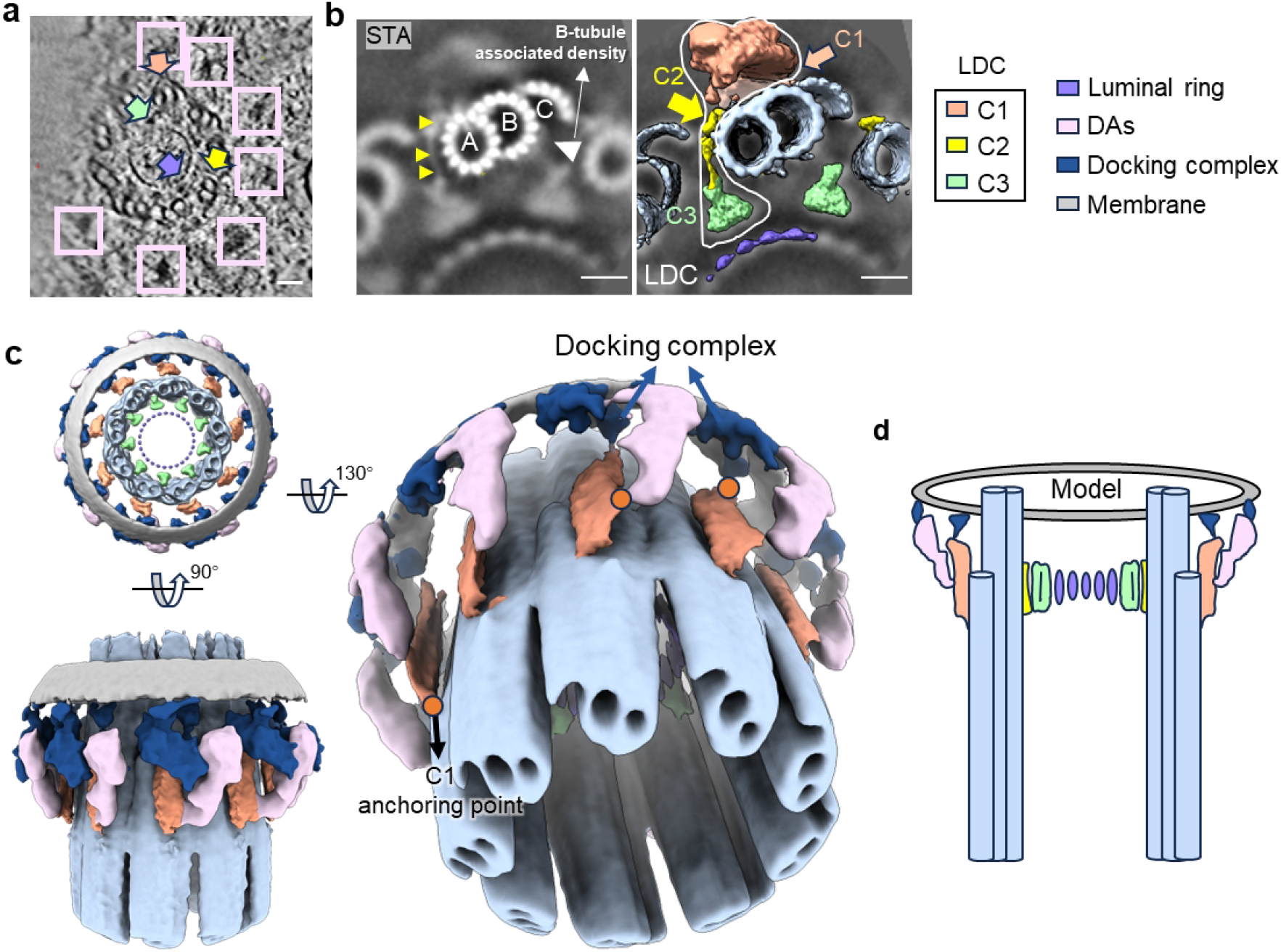
A continuous luminal ring–docking complex links the BB lumen to the plasma membrane. **(a)** Representative tomogram slice of BBs showing luminal ring and DAs connections. Purple arrows, luminal ring; pink rectangles, DAs; orange arrows, C1 of LDC; yellow arrows, C2 of LDC; green arrows, C3 of LDC. Scale bar: 50 nm. **(b)** STA map of luminal ring showing LDC densities. White arrowhead indicates B-tubule associated density which connects to C2 and C3 of LDC. Scale bar: 50 nm. **(c)** Different views of the map showing the continuous density between luminal ring and membrane. **(d)** Proposed model for a continuous structural pathway connecting the luminal ring, docking complex and plasma membrane.

We identified a previously uncharacterized connector between the luminal ring and distal appendages, which we term the Luminal ring-Distal appendages Connector (LDC) (**Fig. 4b-c**). The LDC comprises three distinct densities (C1-C3), with C2CD3 likely contributing to this “inside-to outside” architecture linkage^29^. Specifically, C1 extended toward the outer wall of A- and B-C out junction (CEP78) in CCd region and contacted C2, which emerges from the A-tubule outer wall. C2 then links to C3, which projects inward to reach the luminal ring (**Fig. 4c**).

Notably, we detected an additional contiguous density that linked both the LDC and the DA to the plasma membrane, which we designate the docking complex (**Fig. 4c**). Together, these modules form a continuous structure spanning from the luminal ring, through the LDC and DA, to the docking complex and ultimately the plasma membrane (**Fig. 4d**). This assembly reinforce BB anchorage at the cell surface and preserves membrane attachment during ciliary beating. This observations are consistent with the established role of DAs in BB docking during ciliogenesis^30^.

Consequently, the CCd region appears to adopt an incomplete C-tubule configuration as a compromise between mechanical continuity and spatial remodelling. Retaining part of the C-tubule connects the outer densities of C1 (**Fig. 4b-c**) and B-tubule associated densities that connect to C2 and C3 (**Fig. 4b**), whereas a complete C-tubule would likely block the space required for LDC passage or insertion. Thus, half-C tubules may serve as a transitional assembly that maintains outer mechanic support while relieving steric constraints during the conversion from a BB MTT architecture to a distal MTD-based architecture.

Together, our results define a continuous structural conduit from the BB lumen to the plasma membrane, indicating that the luminal ring is directly integrated into the membrane-anchoring interface. This organization coordinates BB structural stability, membrane docking and spatial positioning during motile ciliogenesis.

### Augmin binds GCP4 to anchor luminal γ-TuRC for CP microtubules biogenesis

γ-TuRC has recently been identified within the centriole lumen, where they are anchored to the centriolar microtubule wall and contributes to spindle apparatus assembly^13,31–33^. Here, we found that BBs also harbour abundant luminal γ-TuRCs (> 40 per BB), arranged in a regular array ∼45 nm from the MTTs (**Fig. S21 and Fig. S22a-b**). A prominent feature in the tomogram is a distinct linker density connecting γ-TuRC to the MTT (**Fig. S22c-d**), which resembles the augmin complex in both morphology and dimensions^33,34^. These observations suggest that luminal γ-TuRCs are anchored within the BB via augmin, consistent with previous reported in centrioles^6,33^.

To further dissect the *in-situ* structure of the γ-TuRC, we purified γ-TuRCs from mouse brain and determined its high-resolution cryo-EM single-particle analysis structure (**Fig. S22e-f, Fig. S23 and Table S3**). The isolated γ-TuRC predominantly adopted an open conformation. Fitting this model into the *in situ* luminal γ-TuRC map revealed two hook-like densities extending from GCP4, as well as a quadrilateral density at the base of complex (**Fig. 5a**). This quadrilateral basal density has previously been assigned to NEDD1, a proposed adaptor for augmin recruitment. However, we did not observe any additional density tethered to NEDD1 (**Fig. 5a and Fig. S24a-b**). By contrast, the two hook-like densities adjacent to GCP4 were optimally positioned to form the augmin-γ-TuRC interface (**Fig. 5a and Fig. S24a-b**), consistent with observations of γ-TuRC within centriole obtained by ELI-TriScope^28^ in this study (**Fig. S25 and Fig. S26**). AlphaFold2 modelling, together with in vivo co-immunoprecipitation and in vitro surface plasmon resonance assays, further supported a direct interaction between GCP4 to HAUS1 and HAUS4 (**Fig. 5c, Fig. 24c and Fig. S27**).

**Figure 5.**
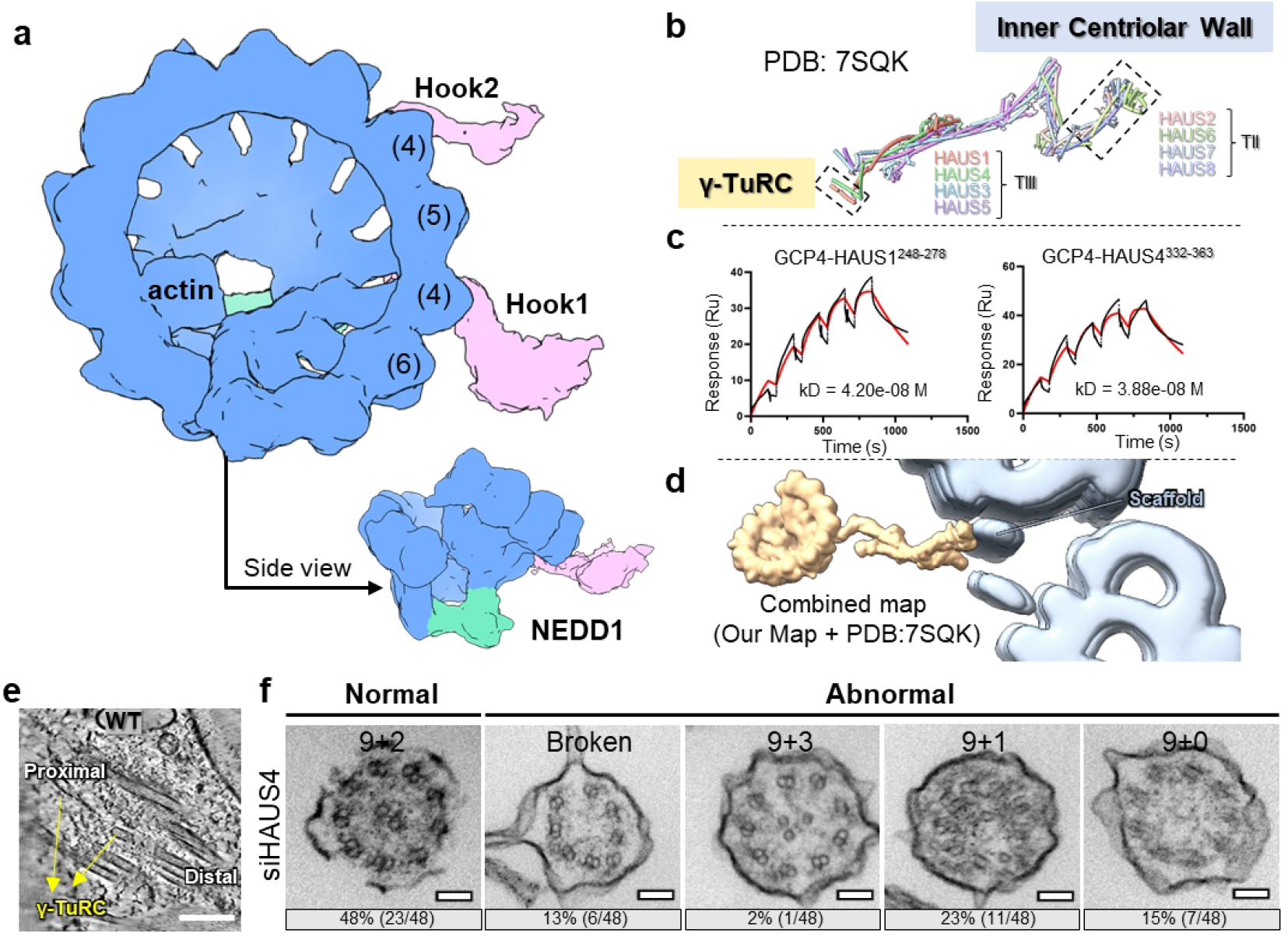
Augmin mediates luminal γ-TuRC anchoring to the BB microtubule wall via GCP4, enabling ciliary CP microtubule nucleation. **(a)** In situ structure of luminal γ-TuRC. The extra densities are indicated and labelled. (b) Component organization of the previously reported augmin structure. (c) In vitro SPR analysis showing direct binding between GCP4 and the C-terminal regions of HAUS1 and HAUS4. (d) The manually assembled γ-TuRC-augmin structure connects well to the BB scaffold. The γ-TuRC-augmin structure was plotted back into the tomogram using the refined coordinates of the MTT and γ-TuRC. (e) Tomogram slice shows luminal nucleated γ-TuRC, also shown in Fig. S28a. (f) TEM images of cilia in siHAUS4 mEPCs. Typical structural defects are shown, together with the number and proportion of each type. Scale bar: 50 nm.

In at least six tomograms, we observed short microtubules emanating from luminal γ-TuRCs, demonstrating that these γ-TuRCs possess microtubule nucleation activity (**Fig. 5e and Fig. S28a**). Notably, previous study has reported that γ-tubulin depletion disrupts ciliary CP assembly^10^, suggesting that CP biogenesis relies on the nucleation activity of luminal γ-TuRCs within the BBs. We quantified the nucleated luminal γ-TuRC across BBs and found that the number of nucleated luminal γ-TuRC never exceeded two, with the corresponding microtubules consistently oriented toward the distal end of BB (**Fig. 5e and Fig. S28a**). These observations support the model that luminal γ-TuRCs serve as nucleation seeds for CP biogenesis in motile cilia.

We further performed RNAi-mediated knockdown against the augmin subunits HAUS1 or HAUS4, and analysed the resulting phenotypes using cryo-ET and IF assay (**Fig. S28-29**). More than 30% of BBs exhibited structural disruption; the ciliary CP was deformed and luminal γ-TuRC was mislocalized (**Fig. S28c and Fig. S29**). 52% of the axonemes no longer possessed the canonical “9+2” architecture and exhibited varying degrees of CP abnormalities, including broken “9+0” (13%), “9+3” (2%), “9+1” (23%) and “9+0” (15%) (**Fig. 5f**). In parallel, cilia length decreased more than 50%, and mislocalization of luminal γ-TuRC rose to 25–50% (**Fig. S29**). Collectively, these observations indicate that loss of the anchoring components disrupts the localization of luminal γ-TuRC, perturbs its arrangement within the BB lumen, and impairs CP biogenesis, leading to abnormal ciliary architecture. Furthermore, a pair of tubular densities located beneath the luminal ring (**Fig. S30a-b**), which were absent in tomograms lacking nucleated microtubules, may be involved in the translocation of nascent microtubule. Therefore, luminal γ-TuRCs and augmin may be involved in microtubule nucleation and nascent central pair microtubules generation in basal body.

## Discussion

### Regionalized architecture of the BBs

BBs are centriole-derived organelles that serve as both the structural template and membrane anchor for cilia. Here, our subnanometer-resolution in situ structural analysis establishes a comprehensive molecular blueprint of the mammalian BB and reveals a fundamental organizational paradigm: spatially confined assembly of CMAPs orchestrates the stepwise proximal-to-distal transitions from MTTs into axonemal MTDs. In detail, proximal CMAPs including RTTN, CEP44 and WDR67 mechanically stabilize MTT architecture and functionally couple the cartwheel-pinhead axis to inter-triplet cohesion. In the central core region, WDR90 and CEP78 act specifically of maintain the integrity of full and half C-tubules, respectively. At the distal BB and TZ, CFAP20 and CAPSL serve as molecular signatures that demarcate the structural transition towards axonemal configuration. Radially, we uncover a continuous inward-to-outward structural conduit spanning from the luminal ring to the plasma membrane. This structural pathway mechanically stabilizes BB anchorage, sustains mechanical tension produced by rhythmic ciliary beating, and achieves functional integration between the internal luminal scaffold and plasma membrane attachment.

Collectively, the spatially compartmentalized organization of BBs underpins its core biological roles in structural maturation, ciliary templating, and persistent membrane anchorage. Building on the structural framework of BBs, we further elucidate a mechanistic model explaining how disease-related CMAP mutations trigger ciliopathies. Such mutations may compromise the stability of regional BB components or disrupting key molecular interfaces indispensable for proper BB assembly (**Fig. S31 and Table S4**). Our findings therefore provide an integrated structural paradigm for interpreting the molecular basis of motile ciliopathies and guiding future mechanistic and therapeutic studies. Notably, the overall architecture of BBs is highly conserved with canonical centrioles (**Fig. S32-33 and Table S5**), reflecting a shared structural framework for microtubule organization. Accordingly, the molecular blueprint of the BB not only advances our mechanistic understanding of ciliary assembly but also provides a critical structural reference for centriole architecture and functional regulation.

### Potential mechanism of ciliary CP biogenesis

Our structural analysis provides insights into the longstanding mystery of ciliary CP biogenesis, suggesting that a spatially and temporally controlled luminal program orchestrated by augmin, γ-TuRC and the luminal ring. Augmin functions as a spatial and geometric regulator: it not only tethers γ-TuRC to the BB scaffold but also ensures the generation of microtubules with correct polarity, using its evolutionary conserved role in directing microtubule nucleation^35^. Depletion of augmin disrupts this tightly regulated cascade: reduced γ-TuRC retention in the BB lumen impairs nucleation efficiency and spatial precision, leading to abnormal “9+0”, “9+1”, or rare “9+3” axonemal patterns. Insufficient augmin shortens the productive nucleation window, causing abortive CP assembly (“9+0” or “9+1”). Furthermore, the failure to initiate timely CP assembly abolishes the feedback signal that restricts additional microtubule entry, resulting in the “9+3” phenotype. Notably, the “9+3” phenotype reflects three uncoordinated single microtubules, rather than one intact CP plus an extra single microtubule. Together, these findings establish a unified structural mechanism for CP biogenesis that links the BB luminal machinery to axonemal patterning, offering a mechanistic framework for understanding ciliopathies arising from defective CP assembly.

## Acknowledgements

We thank Dr. Chuyu Fang and Juyuan Liu from Prof. Xueliang Zhu’s lab for guidance on mEPC culture and BB cluster isolation. We thank all members of Prof. Fei Sun’s lab especially Dr. Yujia Zhai for the standardized management of the laboratory. We thank Dr. Xiaojun Huang, Yan Zeng, Xujing Li, Shuoguo Li, Xixia Li, Lu Qin, Xiaoyun Zhang and Zhongshuang Lv from Centre for Biological Imaging, Interdisciplinary Centre for Biointelligence, Institute of Biophysics, Chinese Academy of Sciences (IBP, CAS), National Multi-mode Trans-scale Biomedical Imaging Centre for assistance in cryo-FIB/cryo-ET and TEM sample preparation. We thank Yuanyuan Chen (IBP, CAS) for SPR experiments, and Dr. Jifeng Wang (IBP, CAS) for MS detection and analysis.

This work was equally supported by grants from National Natural Science Foundation of China (32525005 to F.S., 32471244 to Y.Z.), National Key Research and Development Program (2021YFA1301500 to FS and to 2024YFA1307402 to YZ), Basic Research Program Based on Major Scientific Infrastructures, CAS (JZHKYPT-2021-05 to F.S.), Beijing Natural Science Foundation (JQ24056 to Y.Z.).

## Author contributions

F.S, Y.Z. and X.Z. designed and coordinated this research; Z.W., Y.Z. resolved the structures and wrote the manuscripts; Z.W. and H.Y. performed the biochemical experiments; Z.W. and H.L. prepared the lamellae; Z.W., Y.Z. and F.S. revised the manuscripts.

